# Fast or slow? Clock readout sets internal periods

**DOI:** 10.1101/2023.03.17.533212

**Authors:** Caroline M. Holmes, Stephanie E. Palmer

## Abstract

Circadian clocks are essential for the function of a wide array of organisms, from cyanobacteria to humans. Despite decades of productive study of these systems, mysteries remain, including why many biological clocks have non-24-hour internal periods. We take a new approach to circadian clocks by focusing on downstream readout of the clock’s state to answer physiologically relevant questions, such as ‘when will the sun rise?’ Using this framework, we show that systematic errors arising from sunrise and sunset prediction can be compensated by having non-24 hour internal periods. We show that this prediction holds in models of cyanobacterial circadian clocks. Finally, we predict latitude-dependant qualitative changes in circadian clock structure and the performance of different clock phenotypes in common laboratory experimental setups.

## Introduction

Organisms rely on external sources of energy and metabolites. The availability of these resources often oscillates with a 24-hour period, which means that these fluctuations can be anticipated. This prediction is commonly performed through circadian clocks, which are internal oscillators widely present across the tree of life. These oscillators entrain to external signals, and are read out to implement behavioral programs, such as insulin secretion (1). Significant work over the past several decades has characterized the mechanisms of circadian clocks in diverse organisms from humans and cyanobacteria, and has unified them behind simple, generalizable measurements, such as internal period and phase response curves(2–4).

When an organism with a circadian clock is placed in a constant light environment, it will continue to cycle with a period close to 24 hours. However, these ‘free running’ periods are not *precisely* 24 hours, and vary at least from 23 hours in mice (5) to approximately 24.5 or 25 hours in the plant *Arabidopsis* (6). This period variation does not appear to be an accident of evolutionary noise, given striking patterns in the occurrence of long and short periods. Across even distantly related taxa, nocturnal animals have short periods while diurnal animals often have longer ones (5, 7, 8). This suggests that there may be a functional reason for these differences. Within species, patterns can also be found, such as latitude-dependant variation in the period of clocks in *Arabidopsis* (9, 10). A number of explanations for non-24 hour periods have been posited over the years, including that long or short periods allow for better recovery from phase-slipping errors, or that non-24 hour periods compensate for the differing seasonal timings of dawn and dusk (11). While sensible, these ideas remain largely untested, and do not make clear, experimentally verifiable predictions.

We propose a simple framework to study circadian clock readout in more depth. We assume that the organism only has access to information about time through the clock. In other words, in order to make behavioral decisions related to timing, it must ‘read’ the clock state out. Thus, the circadian clock is an encoder of the physiologically relevant part of the external luminance signal, which is read out downstream. These ‘relevant’ variables are what matters to the particular organism. For example, *Arabidopsis* and cyanobacteria circadian clocks drive expression of a large portion of the genome to a peak that occurs shortly before dawn or dusk (12, 13), suggesting sunrise or sunset timing as relevant variables. *Arabidopsis* uses its clock for a variety of other purposes, including protection from day-active insects (14) and to move their leaf orientation over the course of a day (15). Similarly, cyanobacteria make use of their circadian clock to drive a variety of functions (16–20).

Our framework, which explicitly takes into account the prediction aspect of circadian clocks, explains broad patterns in circadian clock structure, such as non-24 hour internal periods. We focus on cyanobacteria, because its relatively well-understood post translational circadian clock allows for simpler modeling (21), and the relatively short generation times allow for fitness experiments (22). However, our framework is general and can relate readout goals to internal period predictions across species.

## Results

### Clocks as an encoding problem

In our framework, we think of circadian clocks as comprised of two general processes. First, there is an encoding process: the circadian clock changes in time according to its dynamics, and does so in a way that is affected by input light signals. Then, a downstream network must read out the state of the circadian clock in order to answer physiologically relevant questions for the organism (Figure 1a), such as what time it is, or when the sun will rise. Many cyclic behaviors are tied to this kind of information. For example, in cyanobacteria, much of the cell’s behaviors are temporally linked to sunrise or sunset.

**Fig. 1.**
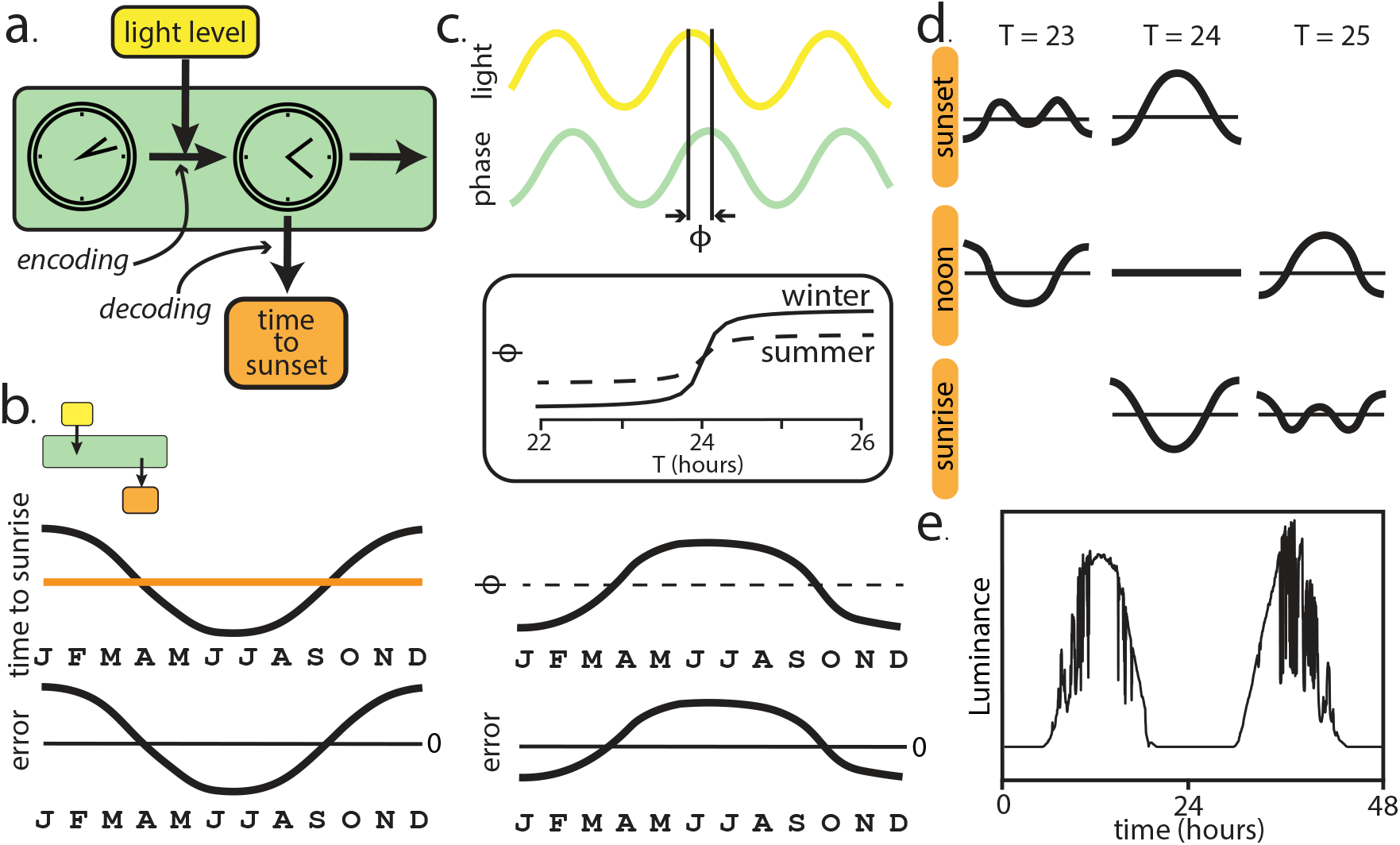
Modeling decoding predicts and relate different systematic errors. a. A cartoon of our framework for circadian clocks. The clock is a dynamical process, which updates at each timestep in a signal-dependent way. This creates an internal state (an ‘encoding’), which then must be read out (‘decoding’) in order to answer physiologically relevant questions. b. One significant pattern in errors relates to sunrise and sunset timings. Here, we show a cartoon: if an organism estimates the time of sunrise by looking at a traditional clock, it will make errors over the course of the year because it will guess the same timing of sunrise every day, even as the sunrise timing changes in a systematic way. c. Non-24 hour periods introduce a different error pattern. We can define a phase lag *φ* as the phase difference between a point in the signal and the a peak in the oscillator. This phase lag will generally be period-dependent (longer periods lag more), and that period-dependence is itself dependant on the amount of driving force set by the brightness of the external light signal. Over the coarse of a year, the amount of driving changes, and this creates a seasonal-varying error. d. A cartoon showing how the errors combine: sunrise and sunset are symmetric problems, as are long and short periods. This means that we expect short periods and sunset prediction to produce low errors, as well as long periods and sunrise prediction. e. All of this is dependent on the structure in the inputs. Rather than *modeling* weather, we use natural light patterns available from NOAA.

We focus primarily on sunrise/sunset and light-level prediction, as these cases illustrate significantly different problems that circadian clocks may aim to solve, and result in different parameter trade-offs. Both of these have plausible physiological relevance for real systems. In cyanobacteria, photoprotection and photosynthesis are important in a way that scales with light level (23, 24), and sunrise and sunset mark the timings of spikes in gene expression (13, 25–27). It is possible that the system cares about other factors, but these can often be categorized as time-like, sunrise/sunset-like, or light-like. Each of these kinds of readout generates systematic errors arising from varying phase offsets.

There are a diversity of biochemical mechanisms for circadian clocks, but in general we can think of clocks as dynamical processes with some periodic structure, which combine their own internal dynamics with external input. These notions can be captured by two general parameters: the gain (how much the system weights the external signal) and the period of the clock. We then can consider how these different parameters (both of which change the nature of the *encoding* in Figure 1a) affect performance at prediction of salient tasks, once the encoding is read out.

If the system is predicting sunrise, one strong, relevant feature is the seasonal variation in the length of the day. Without knowledge of the current seasonal phase, a clock will make systematic errors in estimating dawn. To illustrate this, imagine you are teleported to a random city in the northern hemisphere during the night, sequestered away in a hotel room and asked to predict the time until sunrise. You glance at the bedside alarm clock and see ‘4 A.M.,’ and guess that sunrise is two hours away. You are making a good guess on average, independent of your latitude and season, but in the winter sunrise will generally be more than two hours away (and longer if you are in, say, northern Michigan), and in the summer sunrise is less than three hours away. No matter how accurate your clock is, you will always make seasonal errors. You need other information to make an accurate guess. In Fig. 1b, we illustrate this effect for estimating sunrise with an internal clock.

Having an internal clock with a non-24-hour period also leads to systematic errors in estimating daily luminance landmarks like sunrise. If the internal period is too far from 24-hours, there is constant phase slipping, and the clock is not effective at predicting 24-hour signals. However, there is a range of internal periods which can be entrained to a 24-hour driving signal (28, 29). Within this range, the relative phase of an oscillator and its external driving force varies as a function of period (Fig. 1c). In general, oscillators that are entrained to the signal with long internal periods (*T* > 24 hrs) have a greater phase lag than frequency-matched oscillators, while those with short periods have smaller phase lags. A phase lag in and of itself is not a source of systematic errors. If a clock is always one hour slow, one could look at the time, see ‘5 A.M.,’ and know that it is in fact ‘6 A.M.’ However, the amount of the phase lag is affected not only by the internal period, but also by how strongly the clock is driven by the external light signal. When there is more driving by the input, long and short clocks behave more like 24-hour clocks. This means that for a clock with a long period, the phase lag is high in the winter and high (but smaller) in the summer. This is a clock that is three hours slow in the winter and one hour slow in the summer - when it says ‘5 A.M.,’ one cannot tell whether it is actually winter and ‘8 A.M.,’ or summer and ‘6 A.M.’ without additional information. This lag pattern translates into a season-dependent error, with long periods making errors that are nearly opposite to those introduced by sunrise (Fig. 1c).

If we combine these two kinds of errors, we expect them to counteract each other. Short-period sunset-predictors reduce these systematic errors, as would long-period sunrise-predictors (Fig. 1d). This analysis is based on the interplay between the structure of the input signal and the structure of the clock. In a laboratory setting, organisms are usually presented with simple binary light signals (17, 22, 30–32). However, in the natural environment, sunlight levels rise and decline less abruptly, with variations caused by cloud cover. To provide a more natural input to our model, we use real light traces that were collected by NOAA (Fig. 1e and Methods).

All of the above intuition is reliant on a particular, simplified model of a clock, one where all information must pass through one internal clock readout variable. Real systems have access to other information about time, either through other information streams (i.e., temperature sensors) or through more complicated time-keeping. For example, in cyanobacteria, the circadian clock can be driven by processes dependant on light level (32), as well as by temperature fluctuations (33). Despite the inherent complexity of real biological systems, the dominant features of circadian clocks have been captured by low-dimensional oscillator models, and this means that the intuition discussed above still holds.

### Biologically relevant models

The quantitative effects of these ideas depend on the dynamical system at the heart of the circadian clock as well as the organism’s decoding structure. In order to connect to real biological function, we focus on cyanobacteria, a system with a well-understood, experimentally accessible circadian clock (16, 30, 31, 34, 35).

Unlike mouse, fly, and *Arabidopsis* circadian clocks, which are mediated by gene-expression changes (36, 37), cyanobacterial circadian clocks are fully post-translational (38), which simplifies modeling. This circadian clock, also referred to as the KaiABC system, is made up of a complex of three Kai proteins, which phosphorylate and dephosphorylate in an ordered manner (21, 39). Due to its post-translational nature, this protein-based clock system can be recapitulated *in vitro* with purified proteins (40). Notably, the copy number of these proteins can be very high (as it is in *S. elongatus*), as much as 10, 000 (41), which means that the organism is investing heavily in the circadian clock infrastructure. These *in vitro* oscillators are useful for building biologically relevant models, and make experimental tests of new theories feasible.

This cyanobacterial circadian clock is incredibly important to the function of the organism. Cyanobacteria use their clock to time a wide array of behaviors, from gene expression, which peaks for different genes at dawn or dusk (13, 25–27), to timing nitrogen fixation at night (16), to ensuring that they will have enough nutrients for the night by storing glycogen (17), and timing cell division (18–20). The processes involved in each of these tasks are different, but all of them involve reading out the Kai system to extract information about a few relevant variables. Most of these phenomena are related to sunrise or sunset timing.

Inspired by particular experimental results in the cyanobacterial Kai system (42), we will define a model similar to the one proposed by a recent paper (43). These experiments tracked the limit cycle of KaiC phosphorylation in day-like and night-like conditions *in vitro*, and found that the dominant effect seemed to be movement of the center of a limit cycle.

Here, we will model the circadian clock as a dynamical system,

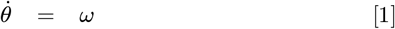

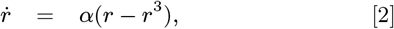

where the origin moves in response to external drive, 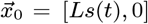, where *L* is a gain parameter and *s*(*t*) is the light signal. We also inject noise at each time step, although all analyses here are done in the low-noise limit, where internal noise is orders of magnitude smaller than the radius of the limit cycle. See Methods for simulation details.

Entrainment by the external light signal happens by moving the origin of the limit cycle. We sketch a cartoon of this model’s behavior in Fig. 2a. If the state traced out exactly the same trajectory in the two-dimensional space every day, the logic from 1 would hold exactly, that seasonal errors resulting from non-24 hour periods and from variation in sunrise timing. Instead, this system has different trajectories on different days, and so the intuition we developed in Fig. 1 should only approximately hold. Numerical simulations are necessary to fully explore and quantify the effects of these interactions.

**Fig. 2.**
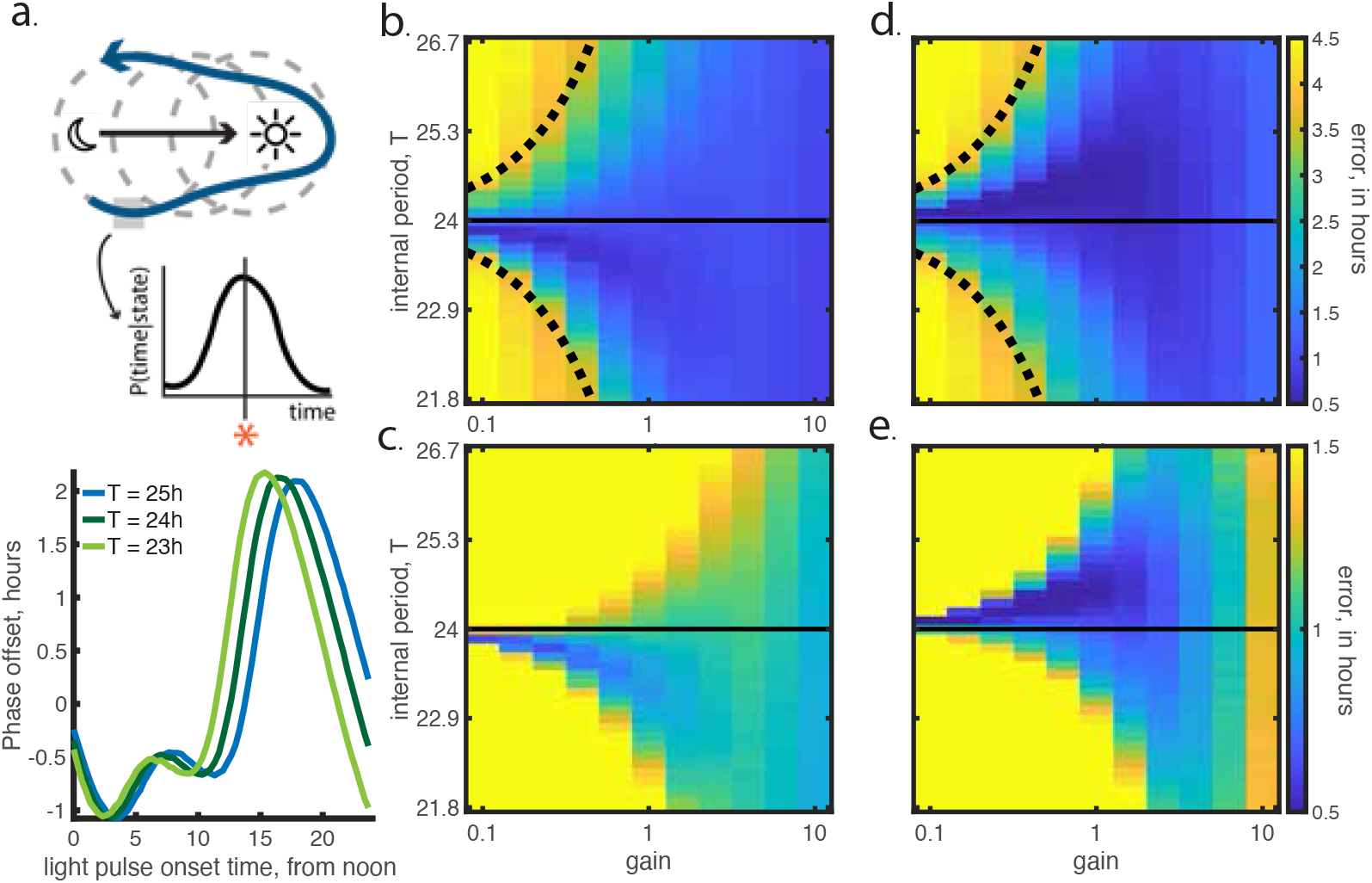
Phenomenological models of cyanobacterial circadian clocks show optimal non-24 hour periods. a. (top) A cartoon of our model of a circadian clock, which is based on experiments in cyanobacteria. We define the decoding of the clock state by discretizing the state and taking the maximum likelihood of the predicted variable. (bottom) Phase response curves, for *L* = 1 and changing period. This shows the response of cells in constant darkness to a pulse of bright light. b. Error in estimating time to sunset, across a range of values for internal period and gain. Dotted line represents boundaries of regime where phase slipping is expected. We find in this model that the optimal period is consistently less than 24 hours, as we had predicted from our Fig. 1. c. Same as b, but with a different color bar to emphasize that the ‘good enough’ region of parameters for clocks is highly asymmetric in period. d, e. Same as b and c, but for sunrise prediction.

We model the decoding process as a maximum likelihood estimate of the external state. We discretize our clock state (*r,θ*) = (*x,y*) ≈ (*x’,y’*). For a system that is using its clock to predict a variable *s*, we have a distribution *P*(*s*|*x’,y’*). For each value of (*x’,y’*) we choose the estimated value of *s* to be the maximum likelihood value of that distribution: 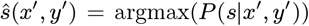. See Methods for details on the decoding.

The cyanobacterial circadian clock does not actually respond directly to light level, but instead can be driven by metabolic processes (44), which increase the ATP/ADP ratio in response to light signals. However, as cyanobacteria photosynthesize, metabolism is a function of light level. Here, we approximate the two signals as a direct link from luminance to readout.

With this framework, we can investigate the relationship between period and what the organism is trying to predict with its clock. In Fig. 2b, the trade-off between gain and internal period for sunset prediction is quantified. For low-gain and large deviations from a 24-hour period (the corners of the plot) we find non-entrainable clocks. These are clocks where the input signal is too weak for the clock to lock to a 24-hour cycle. We plot the boundaries of the 1:1 Arnold tongue as simple Δ*T* = a*L* fits (45). Strikingly, without any parameter tuning, we find that our model gives a similar range of periods compared to those found in real systems, where natural variation in period is on the order of an hour deviation from 24 hours. For example, we can estimate the gain from the experiment that inspired this model as order 1 for *S. elongatus*, and find that periods between 23 and 25 hours entrain well.

As we expected from the logic outlined in Fig. 1, we find that the optimal period is not 24 hours. For sunset prediction (Fig. 2b) the optimal period is significantly shorter than 24 hours, and for sunrise prediction (Fig. 2d) it is consistently longer than 24 hours, as is seen in wild type (WT) cyanobacteria (46). However, we may not expect the real systems to exist at precisely the optimum. Instead, they may occupy some range of nearly-optimal or ‘reasonably good’ solutions, where errors are on the scale of minutes to an hour, but not on the scale of multiple hours. We find that in addition to the optima being located away from 24 hour periods, the regime of good solutions is highly asymmetric (Fig. 2c, e). This suggests that even for near-optimal systems, we expect organisms that care about predicting sunset to have short periods, and those that care about predicting sunrise to have long periods.

### Experimental predictions

Given the dependence of optimal clock structure on the strength of seasonal variation, we can make predictions about the relationship between latitude and period of evolved circadian clocks. As latitude increases, the effects discussed in Fig. 1 increase in magnitude. However, the boundaries of the range of entrainable periods changes as well, as a result of the decreased driving force during the winter months when the external light signal is weaker.

This means that, for a given value of *L*, the range of periods that can effectively be entrained to a 24-hour cycle gets narrower as distance from the equator increases, but also that the strength of the asymmetry increases. This can be seen in Fig. 3a for sunrise prediction, where we used simulated clear-sky environments at each latitude (see Methods). Here, we see that away from the equator, long periods are generally favored, and that the ensemble of clocks that performs within 30 minutes of the optimum becomes more asymmetric with distance from the equator. On the other hand, the optimum approaches 24 hours as a result of the encroachment of the Arnold tongue due to lower driving forces. We note that all simulations in this figure were performed with *L* =1. This matches the order-of-magnitude of the separation between the night and daytime attractors in experiments (35, 42).

**Fig. 3.**
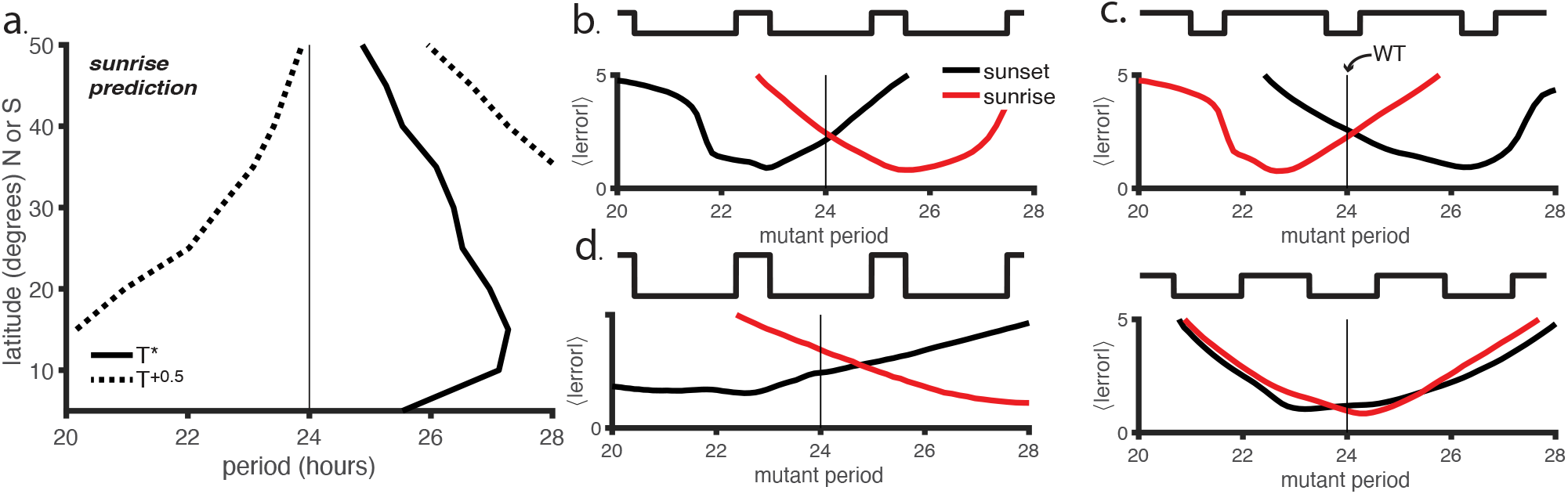
We predict both natural standing variation in circadian clock period, and fitness of mutants that could be measured in laboratory experiments. a. Optimal periods (*T*\*) across latitude (expressed as distance from the equator), for sunrise prediction, and the periods that have half an hour more error than the optimal period (*T*^+0.5^). We find that the optimal period is generally away from 24 hours, as is the region of parameter space that corresponds to nearly-as-good encoding. This translates to a prediction about the distribution of periods that will be found to have evolved at different latitudes. b. Performance of different period mutants to a short photoperiod signal. This corresponds to a prediction about strains that have evolved in the wild with a WT period of 24 hours, and then have had a period mutation in the clock induced in the lab. We find that for organisms where sunrise prediction is important, long mutant periods are favored, and for organisms that predict sunset, short periods are favored. c. Equivalent to b, with a long-photoperiod signal. d. Equivalent to b, with a stronger driving force. e. Equivalent to c, with a 12-hour photoperiod. Here, 24-hour period mutants are optimal, as has been found in prior experimental work.

It is not trivial to make predictions from these results about systems that have evolved at different latitudes, as the optimum and the area around the optimum have different trends. However, we can say that we expect the variance in periods across systems to decrease as latitude increases. Also, except for cases where the period appears exceptionally fine-tuned, we expect the asymmetry of the basin near the optimum to be the dominant factor shaping natural variation in internal periods. We generally expect periods to move away from 24 hours as distance from the equator increases. This has been demonstrated in *Arabidopsis* (10), in *M. guttatas* and soybeans (47), and in insects (48–51), although the trend is not very strong and not entirely consistent. An interesting test of our readout model would be to check the behavior of cyanobacterial circadian clocks from a range of ponds at different latitudes.

We can also make predictions about the performance of cyanobacteria strains with different internal periods in laboratory settings. In cyanobacteria, the fitness of organisms with mutations in their clock genes has been measured in a few light settings (22, 52, 53). These are settings where all of the non-clock machinery evolved in a naturalistic setting, but where the clock is perturbed through mutation by changing some internal parameters, such as the period. Previous work found that in environments where the amount of time with light (photoperiod) equals the amount of time in the dark, matching the period of the clock with that of the stimulus is optimal (22).

In our framework, we can model this effect by fitting the decoder to natural light signals, and then probing behavior in response to different artificial light signals. We find that for a light signal with short photoperiods, the optimal sunset predictor has a period of roughly 23 hours, while the optimal sunrise predictor has a period of roughly 25 hours Fig. 3b. Symmetrically, in a long photo-period environment, the long-period sunset predictors and short-period sunrise predictors are optimal (Fig. 3c).

One may not know if a given organism is a sunrise or sunset predictor, but measuring the competitive of short- and long-period mutants in a short photoperiod environment could answer that question. This would then predict the results of a long-photoperiod experiment. We note that the near-perfect switching of the curves between Fig. 3b and c relies on keeping the amount of driving constant, which is easy to do in our model system. However, even without constant driving, we expect the direction of the optimal period to change with perturbations to the external photoperiod.

Increasing driving (i.e., increasing the external light level) can be expected to stretch our results further period space. Periods even further-from-24-hours should remain ‘fit’ because external drive syncs up the organism to the light environment Fig. 3d. Lastly, we can recapitulate the known results that for equal-photoperiod signals, roughly 24 hour periods are the most fit (22, 52, 53) Fig. 3e.

These predictions are testable for cyanobacteria, and allow for both validation of the theory presented here and for determining the behavioral goal of the circadian clock of any particular cyanobacterial strain.

### Using clocks to predict into the future

We can also consider the problem of prediction explicitly. Our framework assumes that the clock state is encoding a particular variable about the environment, such as the time to sunset, or the light level. However, if acting upon this information takes some amount of time Δ*t*, the encoding should predict the light level at a time Δ*t* in the future Fig. 4a. As some features of the light signal are more predictive than others, Δ*t* will have effects on the optimal encoding.

**Fig. 4.**
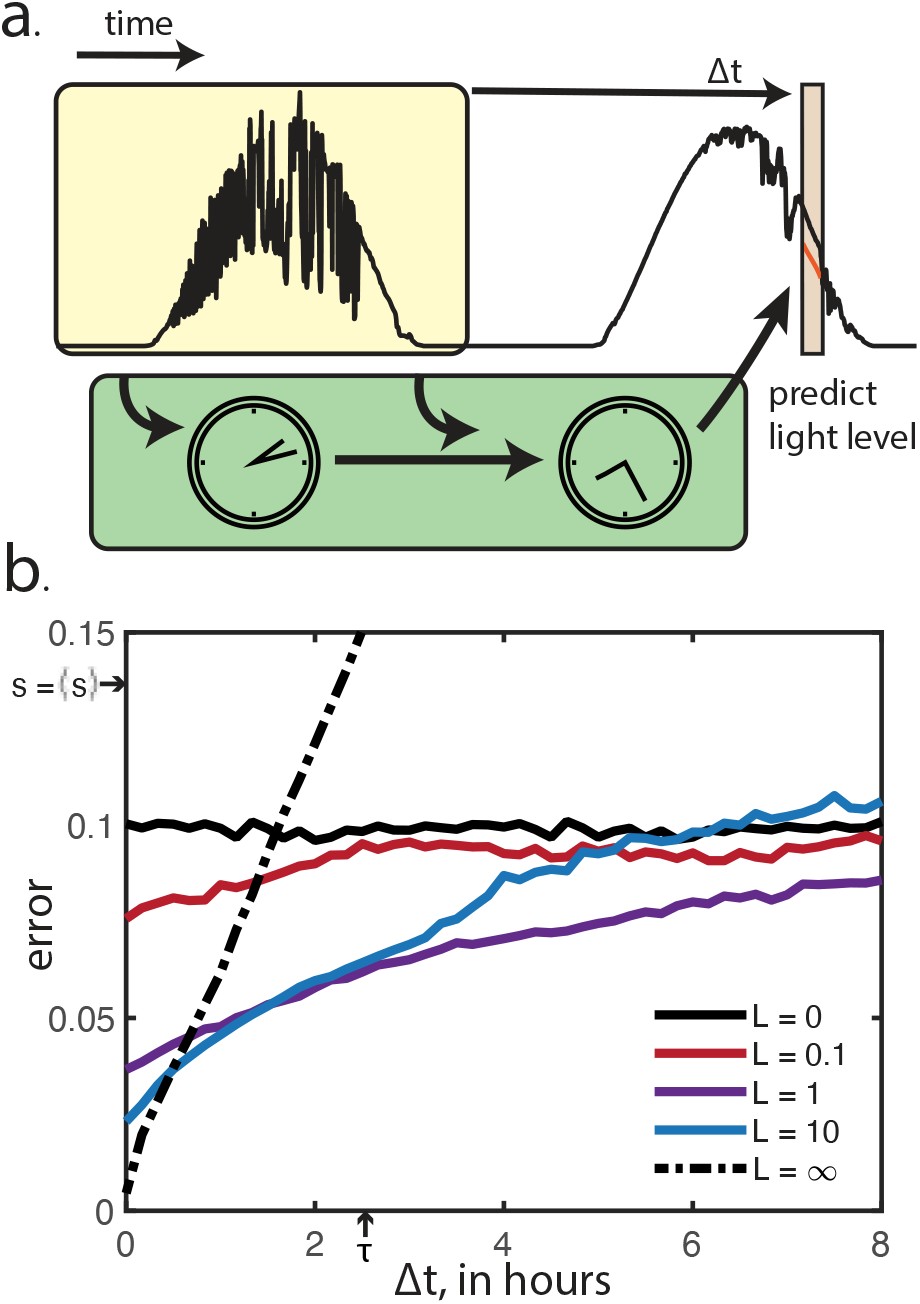
We predict that for time delays of more than half an hour, *L* =1 is optimal. a. We consider the clock as solving a prediction problem, because it is controlling processes which have significant time lags. Because the clock controls a process which takes time Δ*t*, it needs to predict Δ*t* into the future. b. As a function of Δ*t*, we show the performance of different circadian clocks. At very short lags, the sensor (*L* = ∞) is the best predictor, but it is quickly overtaken by intermediateweighting circadian clocks. At no reasonable timescale is a pure clock (*L* = 0) better than a combined sensor. We can compare these timescales to the timescale of autocorrelation in fluctuations in the weather data (*τ*). We can also compare the scale of errors to a purely naive estimate, where the light level is estimated by its average value (*s* = 〈*s*〉).

Here, we will focus on the prediction of light level, rather than on the timing of sunrise and sunset. Just as sunrise and sunset bookend the time in which photosynthesis can be performed, and noon is the time at which maximal energy is available, the light level itself carries information about the quantity of energy available to the organism through photosynthesis (24).

Prior work, at Δ*t* = 0, predicted a monotonic relationship between *L* and performance(43) at light prediction. However, real biological systems have delays at which they act on information.

The main trade-off as Δ*t* increases is not in internal period, but rather in the relative weighting of the current state of the circadian clock and the external signal. This is parameterized by *L* in our system. At the *L* = 0 limit, the system exclusively relies on its own estimates of time, with no external driving, and in the *L* = ∞ limit, the system is entirely externally driven.

At short Δ t, a pure sensor performs well (Fig. 4b), as the current light state carries significantly more information about the near-future light state than is contained in the time. At larger Δ t, as expected, the pure clock outperforms the sensor (dot-dash line). Notably however, for most Δ t’s, intermediate levels of *L*, where both the internal timing and the external signal contribute roughly equally, outperform both the pure clock and the pure sensor. This implies that real organisms likely face a trade-off between these internal parameters that could be revealed by experimental manipulations of the input noise or time delays in the clock readout machinery.

## Discussion

In this paper, we have introduced a new framework for studying circadian clocks, which emphasizes the idea that information about the external environment must be read out effectively by a clock to properly affect behavior. We showed that this framework can explain a long-standing mystery in circadian clocks, that both short and long-period circadian clocks exist throughout the tree of life and have systematic variation with both environmental and behavioral niches. Our results hold quantitatively for models of cyanobacterial circadian clocks, and recapitulate reasonable error values and ranges of entrainable periods. We make concrete predictions about the performance of different-period strains within a particular species. In cyanobacteria, experiments along the lines of those presented in Fig. 3 are doable.

Our framework could be expanded in future work by taking into account multiple information streams. Many circadian clocks can be entrained with multiple signals. For example, cyanobacterial circadian clocks can be entrained with both temperature and luminance signals (33), and this framework could allow investigation of how the two signals are combined.

Our work could also aid in designing new, more informative environmental signals, which could disentangle different answers to the question of what an organism ‘cares about’. For example, key insights may be obscured by probing circadian clocks with binary signals when they have evolved to respond to natural light variation. Our work explicitly inputs real light signals to the model and tests readout errors. This allows for the predictions about the generalizability of different circadian clocks: how well any particular clock performs at latitudes away from where it evolved. This may inform predictions about migration, i.e. about the likelihood of organisms with different clocks migrating within different sets of environments.

Finally, in cyanobacteria, certain species have more sensor- or clock-like circadian responses (41), and our modeling approach could reveal differences between the decoding goals or expected environmental inputs of these different species.

Our results generalize beyond cyanobacteria, and suggest ways to probe the internal sensing goals of other species. Relating sunrise and sunset prediction with period may be challenging to test in longer-generation-time organisms, but larger-scale, systematic analysis of standing variation within populations may reveal clear trends of period with latitude.

By considering circadian clocks as explicitly solving computational readout problems for the organism, we have gained insight into properties of circadian clocks that we expect to generalize across the tree of life.

## Methods

### Light signal

We use natural light data collected by NOAA through the global monitoring library, available at https://gml.noaa.gov/grad/surfrad/index.html. These light intensity signals are collected at several locations within the United States every minute. All analyses shown here were performed on data collected at the Goodwin Creek, MS location. We chose this location because it had some of the largest amounts of data. Specifically, the light signal used here was collected from 1/1/2009-4/7/2021.

We did replicate our analysis on light signals from Boulder, CO, Desert Rock, NV, Fort Peck, MT, Penn State, PA, and Sioux Falls, SD to check for any substantial variation in behavior. We found no change to the qualitative results presented here in the Results.

NOAA provides not only total net radiation measurements (which we use for analysis here), but also several other measurements, including total photosynthetically active radiation. No signal choice that we explored makes a qualitative difference on the results presented. For example, photosynthetically active radiation may be important for cyanobacteria, but this behaves identically to the results we obtain using net radiation.

We did do some minor processing on the light signal. Specifically, NOAA’s sensors occasionally had errors, which are documented. In cases where the error lasted less than 10 minutes, we interpolated the light signal as a straight line between the two boundaries. In cases where the error was more than 10 minutes, we excised that entire day from the light signal, from midnight to midnight.

We measure light levels in units of kWatts/m^2^. In these units, at Goodwin Creek, the average maximum brightness of any day is roughly 0.7, and the maximum brightness observed across all days is about 1.2. This means that units of 1 roughly correspond to the maximal light signal in a day.

For all cases in this paper, we define ‘sunrise’ and ‘sunset’ as when the solar zenith angle crosses 90°.

For simulating light levels at different latitudes, as in Fig. 3a, we followed the calculation in a climatology textbook, (54). We limited our range of latitudes based on data about the geographic range in the oceans (55).

### Simulations: clocks

We ran simulations using the differential equations in the main text, Eqns. 1 2. Our light signal had one-minute time resolution. For our simulations, where Δ*t* < 1 min, we interpolated between these points linearly.

The states of the clocks were initialized randomly, and then were allowed to run for at least five years of light input to converge. This is more convergence time than necessary, but the simulations are fast enough that this was computationally cheap. This allowed for the easy convergence check of whether independent initializations gave the same solution. We then ran the simulations for one additional iteration of the light signal, from which the data were recorded. In cases where internal noise is large, more statistics may be necessary, but all results shown in Results are at low internal noise levels (inspired by the high protein copy number in *S. elongatus* (41)

See the parameter choices section for further simulation details.

### Simulations: decoding

We perform a maximum likelihood decoding. This means that for each ‘clock state,’ we look at the distribution of the readout variable, and define the decoding for that state as the maximum likelihood value of the readout variable.

This requires discretizing the clock state, as the actual state is continuous. Here, we discretize in a 50×50 grid of states, but our results hold qualitatively if this discretization is changed. We checked this by replicating our results for different numbers of discretized states, ranging over an order of magnitude. Our maximum likelihood approach also requires discretizing the readout variable, which is binned into 100 levels for all results shown. Similarly, this choice can be significantly changed without qualitative changes to the results, and we checked this through additional simulations.

For the case of decoding light level, naive applications of maximum likelihood decoding is not effective. Light level has an unusual distribution: for a significant fraction of the day, it is exactly zero, while over the rest of the day it is distributed across a range of continuous values. If we naively discretize the light value, we will find that a light level of zero is nearly always the maximum likelihood choice. To properly perform maximum likelihood decoding, we adaptively bin the light level into equal-probability bins. Many of these bins map to light levels of zero, and we randomly assign zero-light-level points to these bins.

### Parameter choices

For the model described here, every simulation requires the choice of a few parameters: *L, α, T*, the internal noise level, *σ*, and the external light signal, *s*(*t*).

All simulations shown were performed with *α* = 5, but we performed all simulations for both *α* =1.5 and *α* = 5 to check for qualitative effects on our results and found no significant differences.

We additionally show all simulations for a low-noise regime, with *σ* = 10^-3^. Increasing the magnitude of *σ* increases the errors, but does not change our core result that predicts non-24-hour optimal periods.

For Fig. 2a, we entrain our model with 10 days of light/dark cycling. We make use of the artificial light signal created for Fig. 3a, and use the equatorial light signal for this training. We took the light signal to zero at noon on the 10th day, and then we then put the model organism in darkness, except for a one-hour pulse of bright light (brightness = 1). The timing of this pulse is then varied, moving from 0 to 24 hours away from the original stopping out of the light signal. For this figure, periods are as shown, and *L* = 1.

For Fig. 2b-e, we input the natural light data from Goodwin Creek, MS. We use this signal both to train the decoding and to measure performance. *L* and *T* are allowed to vary as shown in the figure.

For Fig. 3a, we set *L* =1. We make this choice as it roughly matches the value of *L* seen in the phosphorylation measurements (35, 42) that inspired our model. We use artificially generated light signals as described above for both training the decoder and measuring performance.

For Fig. 3b-e, we again set *L* = 1. We train the decoder on the same natural light signal used in 2b-e to mimic evolution, with *T* = 24. We then test errors in response to step-wise light signals, with non-24 hour periods. For b, c, and e the light has a magnitude of 0.5 kW/m^2^, while in d it has a magnitude of 1 kW/m^2^. For b, c, d, and e the photoperiods are 8, 16, 8, and 12 hours respectively.

For Fig. 4, we vary *L* as shown, and set *T* = 24.

## ACKNOWLEDGMENTS

We thank Michael Rust, Diane Schnitkey, and Andrew Schober for several useful conversations and comments on this manuscript. This work was supported in part by the National Science Foundation, through the Center for the Physics of Biological Function (PHY-1734030) and a Graduate Research Fellowship (CMH).

## References

1. Sadacca LA, Lamia KA, Delemos AS, Blum B, Weitz C (2011) An intrinsic circadian clock of the pancreas is required for normal insulin release and glucose homeostasis in mice. Diabetologia 54:120–124.

2. Dunlap JC, Loros JJ, DeCoursey PJ (2004) Chronobiology: biological timekeeping. (Sinauer Associates).

3. Feng D, Lazar MA (2012) Clocks, metabolism, and the epigenome. Molecular cell 47(2):158–167.

4. Dunlap JC (1999) Molecular bases for circadian clocks. Cell 96(2):271–290.

5. Daan S, Pittendrigh CS (1976) A functional analysis of circadian pacemakers in nocturnal rodents. Journal of Comparative Physiology A 106(3):253–266.

6. Gould PD, et al. (2006) The molecular basis of temperature compensation in the arabidopsis circadian clock. The Plant Cell 18(5):1177–1187.

7. Aschoff J (1979) Circadian rhythms: influences of internal and external factors on the period measured in constant conditions 1. Zeitschrift für Tierpsychologie 49(3):225–249.

8. Jha PK, Challet E, Kalsbeek A (2015) Circadian rhythms in glucose and lipid metabolism in nocturnal and diurnal mammals. Molecular and cellular endocrinology 418:74–88.

9. Rees H, Joynson R, Brown JK, Hall A (2021) Naturally occurring circadian rhythm variation associated with clock gene loci in swedish arabidopsis accessions. Plant, Cell & Environment 44(3):807–820.

10. Michael TP, et al. (2003) Enhanced fitness conferred by naturally occurring variation in the circadian clock. Science 302(5647):1049–1053.

11. Hirschie Johnson C, Elliott JA, Foster R (2003) Entrainment of circadian programs. Chronobiology international 20(5):741–774.

12. Doherty CJ, Kay SA (2010) Circadian control of global gene expression patterns. Annual review of genetics 44:419–444.

13. Ito H, et al. (2009) Cyanobacterial daily life with kai-based circadian and diurnal genome-wide transcriptional control in synechococcus elongatus. Proceedings of the National Academy of Sciences 106(33):14168–14173.

14. Goodspeed D, Chehab EW, Min-Venditti A, Braam J, Covington MF (2012) Arabidopsis synchronizes jasmonate-mediated defense with insect circadian behavior. Proceedings of the National Academy of Sciences 109(12):4674–4677.

15. Dornbusch T, Michaud O, Xenarios I, Fankhauser C (2014) Differentially phased leaf growth and movements in arabidopsis depend on coordinated circadian and light regulation. The Plant Cell 26(10):3911–3921.

16. Mitsui A, et al. (1986) Strategy by which nitrogen-fixing unicellular cyanobacteria grow photoautotrophically. Nature 323(6090):720–722.

17. Pattanayak GK, Phong C, Rust MJ (2014) Rhythms in energy storage control the ability of the cyanobacterial circadian clock to reset. Current Biology 24(16):1934–1938.

18. Mori T, Binder B, Johnson CH (1996) Circadian gating of cell division in cyanobacteria growing with average doubling times of less than 24 hours. Proceedings of the National Academy of Sciences 93(19):10183–10188.

19. Yang Q, Pando BF, Dong G, Golden SS, van Oudenaarden A (2010) Circadian gating of the cell cycle revealed in single cyanobacterial cells. Science 327(5972):1522–1526.

20. Liao Y Rust MJ (2021) The circadian clock ensures successful dna replication in cyanobacteria. Proceedings of the National Academy of Sciences 118(20):e2022516118.

21. Swan JA, Golden SS, LiWang A, Partch CL (2018) Structure, function, and mechanism of the core circadian clock in cyanobacteria. Journal of Biological Chemistry 293(14):5026–5034.

22. Ouyang Y, Andersson CR, Kondo T, Golden SS, Johnson CH (1998) Resonating circadian clocks enhance fitness in cyanobacteria. Proceedings of the National Academy of Sciences 95(15):8660–8664.

23. Allahverdiyeva Y, Suorsa M, Tikkanen M, Aro EM (2015) Photoprotection of photosystems in fluctuating light intensities. Journal of experimental botany 66(9):2427–2436.

24. Gaastra P (1958) Light energy conversion in field crops in comparison with the photosynthetic efficiency under laboratory conditions. (H. Veenman & Zonen).

25. Markson JS, Piechura JR, Puszynska AM, O’Shea EK (2013) Circadian control of global gene expression by the cyanobacterial master regulator rpaa. Cell 155(6):1396–1408.

26. Vijayan V, Zuzow R, O’Shea EK (2009) Oscillations in supercoiling drive circadian gene expression in cyanobacteria. Proceedings of the National Academy of Sciences 106(52):22564–22568.

27. Liu Y, et al. (1995) Circadian orchestration of gene expression in cyanobacteria. Genes & development 9(12):1469–1478.

28. Strogatz SH (2018) Nonlinear dynamics and chaos with student solutions manual: With applications to physics, biology, chemistry, and engineering. (CRC press).

29. Kuramoto Y, Kuramoto Y (1984) Chemical turbulence. (Springer).

30. Diamond S, Jun D, Rubin BE, Golden SS (2015) The circadian oscillator in synechococcus elongatus controls metabolite partitioning during diurnal growth. Proceedings ofthe National Academy ofSciences 112(15):E1916–E1925.

31. Martins BM, Tooke AK, Thomas P Locke JC (2018) Cell size control driven by the circadian clock and environment in cyanobacteria. Proceedings of the National Academy of Sciences 115(48):E11415–E11424.

32. Rust MJ, Golden SS, O’Shea EK (2011) Light-driven changes in energy metabolism directly entrain the cyanobacterial circadian oscillator. science 331(6014):220–223.

33. Yoshida T, Murayama Y, Ito H, Kageyama H, Kondo T (2009) Nonparametric entrainment of the in vitro circadian phosphorylation rhythm of cyanobacterial kaic by temperature cycle. Proceedings of the National Academy of Sciences 106(5):1648–1653.

34. Ishiura M, et al. (1998) Expression of a gene cluster kaiabc as a circadian feedback process in cyanobacteria. Science 281(5382):1519–1523.

35. Leypunskiy E, et al. (2017) The cyanobacterial circadian clock follows midday in vivo and in vitro. Elife 6:e23539.

36. Allada R, Emery P, Takahashi JS, Rosbash M (2001) Stopping time: the genetics of fly and mouse circadian clocks. Annual review of neuroscience 24(1):1091–1119.

37. Alabadi D, et al. (2001) Reciprocal regulation between toc1 and lhy/cca1 within the arabidopsis circadian clock. Science 293(5531):880–883.

38. Mehra A, Baker CL, Loros JJ, Dunlap JC (2009) Post-translational modifications in circadian rhythms. Trends in biochemical sciences 34(10):483–490.

39. Rust MJ, Markson JS, Lane WS, Fisher DS, O’Shea EK (2007) Ordered phosphorylation governs oscillation of a three-protein circadian clock. Science 318(5851):809–812.

40. Nakajima M, et al. (2005) Reconstitution of circadian oscillation of cyanobacterial kaic phosphorylation in vitro. science 308(5720):414–415.

41. Chew J, Leypunskiy E, Lin J, Murugan A, Rust MJ (2018) High protein copy number is required to suppress stochasticity in the cyanobacterial circadian clock. Nature communications 9(1):1–10.

42. Phong C, Markson JS, Wilhoite CM, Rust MJ (2013) Robust and tunable circadian rhythms from differentially sensitive catalytic domains. Proceedings of the National Academy of Sciences 110(3):1124–1129.

43. Pittayakanchit W, Lu Z, Chew J, Rust MJ, Murugan A (2018) Biophysical clocks face a trade-off between internal and external noise resistance. Elife 7:e37624.

44. Pattanayak G, Rust MJ (2014) The cyanobacterial clock and metabolism. Current opinion in microbiology 18:90–95.

45. Arnold VI (1961) Small denominators. i. mapping the circle onto itself. Izv. Akad. Nauk SSSR Ser. Mat 25(1):21–86.

46. Murayama Y, et al. (2017) Low temperature nullifies the circadian clock in cyanobacteria through hopf bifurcation. Proceedings of the National Academy of Sciences 114(22):5641–5646.

47. Greenham K, et al. (2017) Geographic variation of plant circadian clock function in natural and agricultural settings. Journal of Biological Rhythms 32(1):26–34.

48. Tyukmaeva VI, Salminen TS, Kankare M, Knott KE, Hoikkala A (2011) Adaptation to a seasonally varying environment: a strong latitudinal cline in reproductive diapause combined with high gene flow in drosophila montana. Ecology and Evolution 1(2):160–168.

49. Wang XP, et al. (2012) Geographic variation in photoperiodic diapause induction and diapause intensity in sericinus montelus (lepidoptera: Papilionidae). Insect Science 19(3):295–302.

50. Allemand R, David J (1976) The circadian rhythm of oviposition indrosophila melanogaster: A genetic latitudinal cline in wild populations. Experientia 32(11):1403–1405.

51. Hut RA, Paolucci S, Dor R, Kyriacou CP, Daan S (2013) Latitudinal clines: an evolutionary view on biological rhythms. Proceedings of the Royal Society B: Biological Sciences 280(1765):20130433.

52. Woelfle MA, Ouyang Y, Phanvijhitsiri K, Johnson CH (2004) The adaptive value of circadian clocks: an experimental assessment in cyanobacteria. Current Biology 14(16):1481–1486.

53. Ma P, Woelfle MA, Johnson CH (2013) An evolutionary fitness enhancement conferred by the circadian system in cyanobacteria. Chaos, Solitons & Fractals 50:65–74.

54. Hartmann DL (2015) Global physical climatology. (Newnes) Vol. 103.

55. Flombaum P, et al. (2013) Present and future global distributions of the marine cyanobacteria prochlorococcus and synechococcus. Proceedings of the National Academy of Sciences 110(24):9824–9829.

